# Haplotype-based Parallel PBWT for Biobank Scale Data

**DOI:** 10.1101/2025.02.04.636317

**Authors:** Kecong Tang, Ahsan Sanaullah, Degui Zhi, Shaojie Zhang

**Affiliations:** Department of Computer Science, University of Central Florida, Orlando, FL 32826, USA; McWilliams School of Biomedical Informatics, University of Texas Health Science Center at Houston, Houston, TX 77030, USA

**Keywords:** PBWT, Parallel Computing, Haplotype Matching

## Abstract

Durbin’s positional Burrows-Wheeler transform (PBWT) enables algorithms with the optimal time complexity of *O*(*MN*) for reporting all vs all haplotype matches in a population panel with *M* haplotypes and *N* variant sites. However, even this efficiency may still be too slow when the number of haplotypes reaches millions. To further reduce the run time, in this paper, a parallel version of the PBWT algorithms is introduced for all versus all haplotype matching, which is called HP-PBWT (haplotype-based parallel PBWT). HP-PBWT parallelly executes the PBWT by splitting a haplotype panel into blocks of haplotypes. HP-PBWT algorithms achieve parallelization for PBWT construction, reporting all versus all L-long matches, and reporting all versus all set-maximal matches while maintaining memory efficiency. HP-PBWT has an 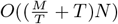 time complexity in PBWT construction, and an 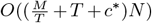 time complexity for reporting all versus all L-long matches and reporting all versus all set-maximal matches, where *T* is the number of threads and *c\** is the maximum number of matches (of length L or maximum divergence value for L-long matches and set-maximal matches, respectively) per haplotype per site. HP-PBWT achieves 4-fold speed-up in UK Biobank genotyping array data with 30 threads in the IO-included benchmarks. When applying HP-PBWT to a dataset of 8 million randomized haplotypes (random binary strings of equal length) in the IO-excluded benchmarks, it can achieve a 22-fold speed-up with 60 cores on the Amazon EC2 server. With further hardware optimization, HP-PBWT is expected to handle billions of haplotypes efficiently.

## 1 Introduction

The positional Burrows–Wheeler transform (PBWT) [6] is an efficient data structure for finding haplotype matches and data compression. Construction of the PBWT can be done in linear time with respect to the number of haplotypes times the number of sites. The original PBWT paper also introduced efficient algorithms for reporting all versus all haplotype matches in a genetic panel. These algorithms have been widely applied in Identity-By-Descent (IBD) segment detection [8, 11, 20], genotype imputation [5, 13], and haplotype phasing [4, 9]. However, once the haplotype dimension of biobank panels exceeds many millions, even the PBWT may not be fast enough to satisfy the need for speed. The rapidly growing total number of genotyped individuals could reach billions in the future. This means an efficient parallel version of PBWT suitable for processing large-scale data input is in high demand.

Wertenbroek et al. [19] presented a parallelized PBWT by dividing the panel into sub panels with ranges of sites, then applied a merging algorithm to generate the final output. Their algorithm has an 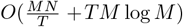 time complexity, where *M* is the number of haplotypes, *N* is the number of sites, and *T* is the number of threads. However, their algorithm needs to load the whole haplotype panel into memory or read the haplotype panel multiple times to perform the dividing and merging steps. Therefore, this approach is best applicable to panels with sequencing data of a relatively small sample size [1, 18].

On the other hand, the haplotype dimension of the panel is the more promising dimension to parallelize. Currently, genotyping array density datasets with millions of haplotypes [10, 16, 17] are widely used in both research and commercial applications. When compared to sequencing datasets, genotyping array density datasets have sparser sites, while typically having many more haplotypes, sometimes millions. Moreover, in the not-too-distant future, the number of genotyped samples could reach billions. Durbin’s algorithms, even though enjoying a linear complexity to the number of haplotypes, need parallelization to scale up further.

In this paper, a haplotype-based parallel PBWT, HP-PBWT, is proposed, which is suitable for processing billions of haplotypes. The HP-PBWT is designed to the break dependencies in the prefix array computation and divergence array computation in Durbin’s original PBWT. Furthermore, two additional efficient algorithms are designed to report all versus all L-long matches and all versus all set-maximal matches using the parallelized PBWT. HP-PBWT is memory efficient, it maintains the efficient sweeping behavior in Durbin’s original PBWT and only needs an *O*(*M*) memory space. HP-PBWT has 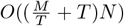 run time for constructing the PBWT, and 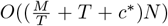 run time for reporting all versus all L-long matches and reporting all versus all set-maximal matches, where *T* is the number of threads, and *c\** is the maximum number of matches per haplotype per site and in practice *c\** ≪ *M*.

## 2 Background

The description in this work follows Durbin’s original work [6]. The initial input of Durbin’s PBWT is *X*, an *M* by *N* two-dimensional binary array, which has *M* binary strings, *x*_*i*_, which each represents a haplotype, and each haplotype has *N* sites. The fundamental outputs are a prefix array *P*_*k*_ and a divergence array *D*_*k*_ during the sorting process for each site 0 ≤ *k < N*.

The prefix array *P*_*k*_ stores haplotype IDs according to the colexicographic order of the haplotype prefixes of length *k* + 1. *P*_*k*_ stores a permutation of [0, *M* − 1] such that 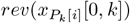 is lexicographically smaller than 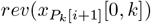 for all *i*, where *rev*(*a*) is the reverse of a string *a*. The term *Y*_*k*_ in Durbin’s PBWT refers to a permutation of all sites in *X* at *k*-th location *X*_*k*_, which is sorted based on the haplotype IDs in *P*_*k−*1_ such that 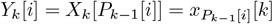.

The divergence array *D*_*k*_ indicates the length of the match at *k* between two adjacent haplotypes in the sorted order of *P*_*k*_. Therefore, if *P*_*k*_[*a*] = *i, P*_*k*_[*a* −1] = *i*′, and *D*_*k*_[*i*] = *j*, then, 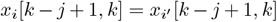 and 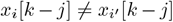. Note that the definition of the divergence array here is slightly different from Durbin’s original definition. Durbin defined the divergence value as storing the starting position of the match and the values were permuted by the prefix array sorting.

A match between sequences *x*_*a*_ and *x*_*b*_ on [*i, j*] is locally maximal if *x*_*a*_[*i, j*] = *x*_*b*_[*i, j*] and it cannot be extended in either direction. I.E. if *x*_*a*_[*i* − 1] ≠ *x*_*b*_[*i* − 1], and *x*_*a*_[*j* + 1] *x*_*b*_[*j* + 1]. Given a length cut-off *L*, haplotypes *x*_*a*_ and *x*_*b*_ have an L-long match on [*i, j*], if [*i, j*] is a locally maximal match and *j* −*i* + 1 ≥*L*. Durbin’s Algorithm 3 outputs all L-long matches between all pairs of haplotypes. In this paper, this is referred to as outputting all versus all L-long matches.

A set-maximal match is defined on a haplotype *x*_*a*_ and a set of haplotypes, *X*. If *x*_*a*_ has a set-maximal match to *x*_*b*_ within *X* on [*i, j*], then there is no larger match that contains it. I.E. ∀*x*_*c*_ ∈ *X, x*_*a*_[*i* − 1, *j*] ≠ *x*_*c*_[*i* − 1, *j*] and *x*_*a*_[*i, j* +1] ≠ *x*_*c*_[*i, j* +1]. Durbin’s Algorithm 4 outputs the set-maximal matches between *x*_*d*_ and *X* \{*x*_*d*_} for all *x*_*d*_ ∈ *X*. In this paper, this is referred to as outputting all versus all set-maximal matches.

## 3 Methods

The focus of this work is to reduce the time complexity of the *M* dimension, the *N* dimension remains the same. First, a parallel prefix sum algorithm is used to compute the prefix array in parallel. Second, *D*_*k*_ is calculated by dividing *Y*_*k*_ into *T* partition blocks with *T* threads independently in parallel. Third, an efficient fine-grained parallel algorithm is designed to report all versus all L-long matches. At the end of this section, all versus all set-maximal matches are also reported in parallel with a similar algorithm. These algorithms Have 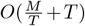 span per site for PBWT construction and 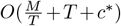 span per site for reporting all versus all L-long matches and reporting all versus all set-maximal matches, where *T* is the number of threads, and *c\** is the maximum number of matches per haplotype per site and in practice *c\** ≪ *M*.

### 3.1 Parallel Prefix Array Computation

Durbin’s Algorithm 1 computes *P*_*k*_ from *P*_*k−*1_ and *Y*_*k*_, by placing *i* ∈*P*_*k−*1_ into a “zero” container if *Y*_*k*_[*i*] = 0 or a “one” container if *Y*_*k*_[*i*] = 1 sequentially. Then, *P*_*k*_ is constructed by concatenating the “zero” and “one” containers. Since the process is done sequentially, this dependency has to be removed in order to execute in parallel.

The *P*_*k*_ construction of Durbin’s PBWT Algorithm 1 is converted into a prefix sum problem. The first step is to run prefix sum on *Y*_*k*_ to create a *PSA*_*k*_ such that 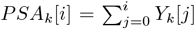. An important property of the *PSA*_*k*_ is, the *PSA*_*k*_ indicates the starting location of “1”s in *P*_*k*_. The *PSA*_*k*_[*M* −1] is the number of “1”s in *Y*_*k*_, and the offset *OS* = *M PSA*_*k*_[*M* −1] is the number of “0”s in *Y*_*k*_. The prefix array is defined as *P*_*k*_ = *PZ*_*k*_ + *PO*_*k*_, where *PZ*_*k*_ is the “zero” container and *PO*_*k*_ is the “one” container. The *PZ*_*k*_ holds all haplotype indices *i* such that *Y*_*k*_[*i*] = 0 by the order in *P*_*k−*1_. Similarly, the *PO*_*k*_ holds the haplotype indices *i* such that *Y*_*k*_[*i*] = 1. A mapping example is introduced from *P*_*k−*1_ to *P*_*k*_ according to *PSA*_*k*_ in Figure 1. The *PO*_*k*_ part of *P*_*k*_ can be populated by *P*_*k*_[*OS* + *PSA*_*k*_[*i*] − 1] = *P*_*k−*1_[*i*]. Since *PSA*_*k*_ only increases by one when *Y*_*k*_[*i*] = 1, the value at *PSA*_*k*_[*i*] for haplotype *P*_*k−*1_[*i*] is the exact one-based index of *P*_*k−*1_[*i*] in *PO*_*k*_. The *PZ*_*K*_ is the first part of *P*_*k*_, the new location of a haplotype *P*_*k*_[*i*] is the old index *i* minus the number “1”s appeared before *Y*_*k*_[*i*] which is *PSA*_*k*_[*i*]. Then *PZ*_*k*_ part of the *P*_*k*_ can be computed by *P*_*k*_[*i PSA*_*k*_[*i*]] = *P*_*k−*1_[*i*]. There is no dependency after creating the *PSA*_*k*_, so *P*_*k*_ can be populated in parallel.

**Fig. 1.**
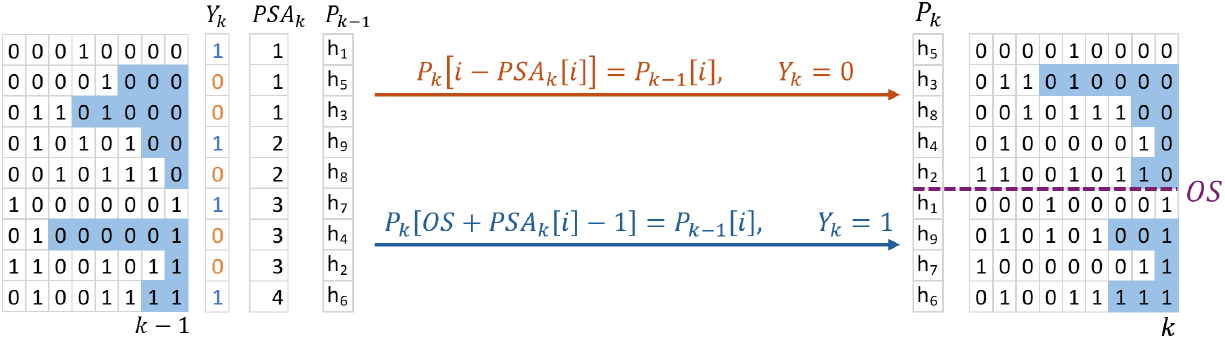
Computation of new prefix array *P*_*k*_ from old prefix array *P*_*k−*1_ in parallel with the help of prefix sum array *PSA*_*k*_. The offset is calculated by *OS* = *M − PSA*_*k*_[*M −* 1] which is the number of zeros in *Y*_*k*_.

Therefore, the computation of the PBWT prefix array has been converted to a prefix sum problem. The prefix sums of *Y*_*k*_ are computed in parallel. First, the input is divided into *T* partition blocks. Then local prefix sums within the partition blocks are calculated using one thread per partition block in parallel. Now each partition block *b* has its locally correct prefix sum values. To acquire the final globally correct prefix sum values, each value in partition block *b* has to be added a proper offset, which is the last prefix sum value of partition block *b* − 1. These offsets need to be computed and added from each partition block sequentially. At the end, the corresponding offset is added to each partition block in parallel. The offset calculation is the only communication stage, and it has an *O*(*T*) time complexity. This version has *O*(*M* + *T*) work and 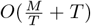 span per site. If *T* is large, then the parallel prefix sum in Section II.E of [2] (referred to as the halving merge algorithm) can be used to reduced the *O*(*T*) term to *O*(log *T*) for a total work of *O*(*M*) and span of 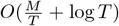 per site. After computing *PSA*_*k*_, the haplotype IDs are mapped to the correct final prefix array location. Each haplotype takes *O*(1) time to map and all *M* haplotypes are processed in parallel (See Appendix A Algorithm 1 for details). Therefore the parallelized prefix array computation algorithm as described has *O*(*M*) work and 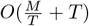 span 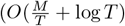 span per site if the algorithm of [2] is used), where *T* is the number of threads, and *T* also equals to the number of partition blocks. The 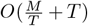 span parallel prefix sum algorithm is used in this work.

### 3.2 Parallel Divergence Array Computation

In Durbin’s algorithm, *D*_*k*_ is computed by sweeping through *P*_*k−*1_ keeping track of the minimum matching lengths of haplotypes that have “0” and “1” in *Y*_*k*_ (*p* and *q* respectively in Durbin’s Algorithm 2) through *M* haplotypes during the process of creating *P*_*k*_. The algorithm checks if a haplotype still matches to its previous upper neighbor haplotype in *P*_*k−*1_ at *Y*_*k*_ site.

On the contrary, once a site is divided into partition blocks, and each block is assigned with one thread, to calculate all the divergence values within a single partition block, the passing down values (*p* and *q* in Durbin’s Algorithm 2) should not be acquired from the thread that assigned to the previous partition block. Otherwise, the whole process becomes sequential. Now the challenge becomes how to acquire the correct passing down divergence values for each partition block in parallel.

Two necessary initial divergence values are calculated for each partition block by searching the previous partition block(s). To further explain, the two divergence values are: the divergence value of the first haplotype in the partition block that has “0” value in the *Y*_*k*_, and the first haplotype in the partition block that has “1” value in the *Y*_*k*_. Call the *n*-th partition block 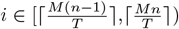, then, all the divergence values of the *n*-th partition block are *D*_*k*_[*P*_*k*_[*i*]]. These two divergence values can be computed by checking some upper *b*-th partition block(s) that *b < n*. The first divergence value is *D*_*k*_[*P*_*k*_[*j*]] such that 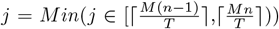 and *Y*_*k*_[*j*] = 0. The second divergence value is *D*_*k*_[*P*_*k*_[*j*]] such that 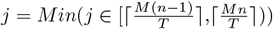 and *Y*_*k*_[*j*] = 1. For any of the two divergence values, *D*_*k*_[*P*_*k*_[*j*]] that *Y*_*k*_[*j*] = *v* and *v* = 0 or 1, *D*_*k*_[*P*_*k*_[*j*]] is computed by: first, find the index *si* that *Y*_*k*_[*si*] = *v, si < j*, and ∀∈*i* (*si, j*) and *Y*_*k*_[*si*] ≠ *v*; then, *D*_*k*_[*P*_*k*_[*j*]] = *Min*(*D*_*k*_[*P*_*k*_[*i*]]) that *i* ∈ (*si, j*]. For each partition block, Algorithm 2 in Appendix A searches the two divergence values in the previous partition block(s). After computing these two divergence values, it can simply loop through the partition block with Durbin’s Algorithm 2 to compute the rest of the divergence values within this partition block in a single thread. The searching step is called upper search. This upper search may cross multiple partition blocks, and it does have a worst case *O*(*M*) time complexity, but this is unlikely to happen since the selected markers in the biobank data panels usually do not have nearly singleton minor allele frequency. Optimizations are applied to prevent the long upper search from happening. The first optimization is verifying if the haplotype *j* is the first “1” in *Y*_*k*_, by checking if *PSA*_*k*_[*P*_*k*_[*j*]] = 1, if so, this divergence value is set to 0, and the upper search isn’t conducted.

The upper search examples are shown in Figure 2 which compute the two passing down divergence values for *D*_*k*_[*h*_9_] in partition block 2 and *D*_*k*_[*h*_12_] in partition block 3. Computing *D*_*k*_[*h*_9_] is simply *D*_*k*_[*h*_9_] = *D*_*k−*1_[*h*_9_] + 1, since the match between haplotype *h*_9_ and *h*_8_ continues. Haplotype *h*_12_ will be sorted after haplotype *h*_1_, so *D*_*k*_[*h*_12_] is minimum divergence value (*D*_*k*_[*h*_3_]) between *h*_12_ and *h*_2_. It has to reach partition block 1 to find the first upper haplotype *h*_1_ that has the same value “1” to haplotype *h*_12_ at *k*-th site.

**Fig. 2.**
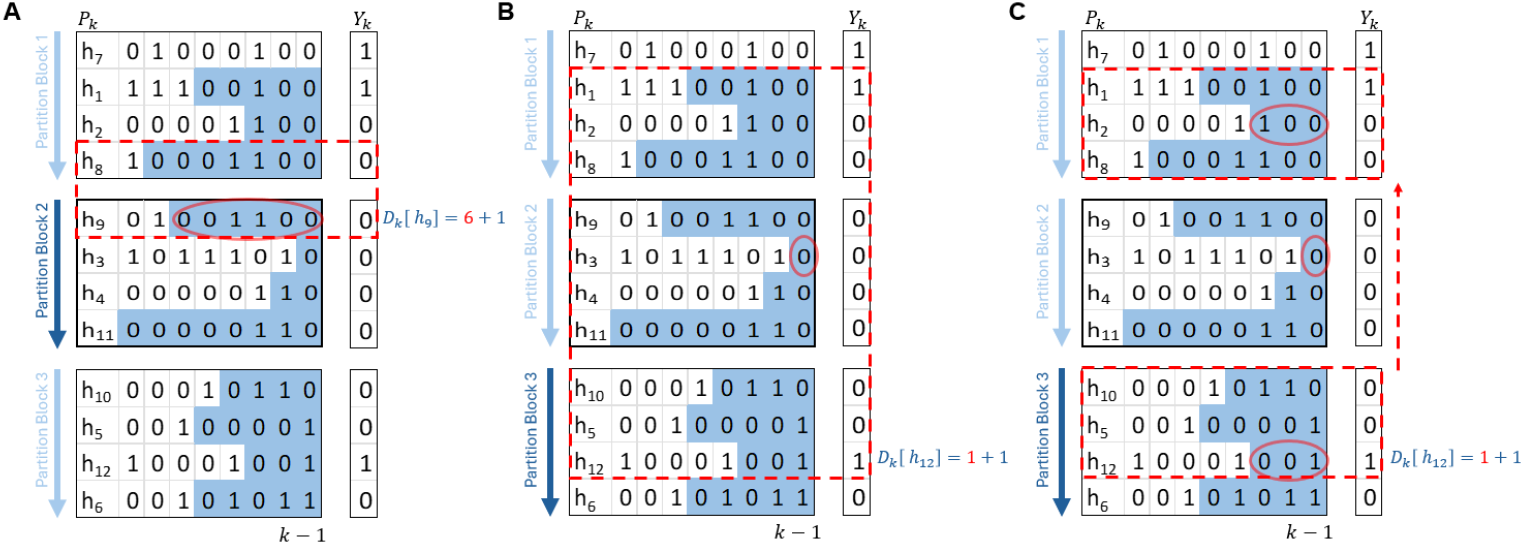
Cross block upper search during divergence value computation. *D*_*k*_[*h*_9_] (in A) needs a short upper search, and *D*_*k*_[*h*_12_] (in B) needs a long upper search that cross block. Additional optimization (in C) by skipping block(s) is designed for the long upper search.

An additional optimization is applied for the situation that there is small amount of “1” in *Y*_*k*_, so the upper search for the haplotype with “1” may end up *O*(*M*). Demonstrated in Figure 2(C), assuming there could be many partition blocks that full of “0”s between *h*_12_ and *h*_1_. For each partition block *B*_*i*_ = [*s, e*] with an individual thread *T*_*i*_, the first step is to identify whether this partition block has “1” or not. Here, a zero-block is defined as if *PSA*_*k*_[*s*− 1] = *PSA*_*k*_[*e* −1], that is to say there are no “1”s in this partition block. If the partition block *B*_*i*_ has “1”, the thread *T*_*i*_ will skip all the zero-block(s) above it and find the nearest upper partition block *B*_*j*_ = [*s*′, *e*′] that has “1” by checking the upper partition blocks according to the *PSA*_*k*_ as shown in Appendix A Algorithm 2. Then thread *T*_*i*_ runs from *e*′ to *s*′ to find the first *p* that *Y*_*k*_[*p*] = 1, and compute the minimum divergence value *minD*_0_ = *D*_*k−*1_[*P*_*k*_[*h*]] that ∀*h*∈ (*p, e*′]. In the meantime, the minimum divergence values for each the skipped zero-blocks are still needed. The minimum divergence values for the zero-blocks are computed by the thread assigned to each partition block once a partition block is identified as zero-block. *T*_*i*_ waits to collect these minimum divergence values for the skipped zero-blocks if they are not yet calculated. Since it is unlikely that a site has a large run of “1”s in practice, in the implementation of HP-PBWT, this optimization is only applied on the “0”s.

The divergence value computation in HP-PBWT has *O*(*M* + *Tr*) work and 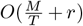 span per site, where *r* is the cost of the upper search and *T* is the number of threads. *r* is reduced to 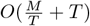 work if the same optimization was applied for both “0”s and “1”s (note *T* ≪ *M*). Then the final work and span of divergence value computation is *O*(*M* + *T*) and 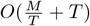 per site respectively.

### 3.3 Parallel Reporting of All versus All L-long matches

Given a set of *P*_*k*_, *D*_*k*_, and *Y*_*k*+1_, Durbin’s Algorithm 3 outputs all matches with length ≥*L* that end at *k*-th site. The algorithm has two basic steps. Step 1, acquire a precise matching block range [*s, e*] that ∀*i* ∈ [*s, e*] *D*_*k*_[*P*_*k*_[*s*]] ≥ *L, D*_*k*_[*P*_*k*_[*e*]] ≥ *L, D*_*k*_[*P*_*k*_[*s* − 1]] *< L*, and *D*_*k*_[*P*_*k*_[*e* + 1]] *< L*. That is to say all pairs of prefixes of haplotypes within the block have longest common suffixes of length at least length *L*. Step 2, output a match between every pair of haplotypes with differing values in *Y*_*k*+1_ within [*s, e*].

In parallel execution, the simplified Algorithm 3 in Appendix A reports matches for each haplotype in an individual thread. For each haplotype *P*_*k*_[*h*], it loops through the haplotypes *P*_*k*_[*i*] ∈ (*h, M*) until *D*_*k*_[*P*_*k*_[*i*]] *< L*, in the meantime it outputs matches if *Y*_*k*+1_[*P*_*k*_[*h*]] ≠ *Y*_*k*+1_[*P*_*k*_[*i*]]. Figure 3 provides an example of thread *I* reports L-long matches for haplotype *h*_4_ independently. In detail, this algorithm does have 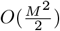 work, since there may be *O*(*M* ^2^) matches at a site. If *c\** is used to represent the maximum number of haplotypes that match with haplotype *i* length *L* or more ending at the current site for all *i*, HP-PBWT has 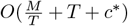 span per site for reporting all versus all L-long matches, where *T* is the number of threads (note that in practice *c\** ≪ *M*).

**Fig. 3.**
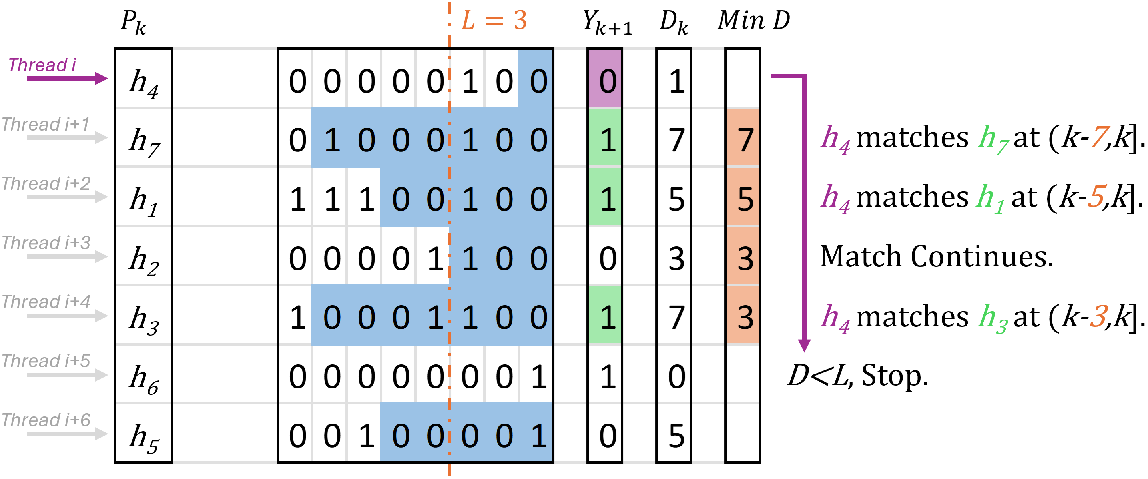
Reporting all versus all L-long matches where *L* = 3 and sorted at *k* site in parallel. *Thread i* reports matches for haplotype *h*_4_, it loops to haplotype *h*_6_ since *D*_*k*_[*h*_6_] *< L*, meanwhile it compares the values of *h*_7_, *h*_1_, *h*_2_ and *h*_3_ in *Y*_*k*+1_ to *h*_4_, then reports the match between *h*_4_ and *h*_7_, the match between *h*_4_ and *h*_1_, and the match between *h*_4_ and *h*_3_.

### 3.4 Parallel Reporting All versus All set-maximal matches

A set-maximal match is a collection of the longest matches ending at *k* site to a single haplotype. [*s, e*] is a range in *P*_*k*_ that contains all the longest matches with haplotype *i* ending at *k*. If a match in [*s, e*] can be extended further, then there are no set-maximal matches ending at *k* for haplotype *i*. For a haplotype *P*_*k*_[*i*], the set-maximal matches at site *k* are reported by: First, compute *D*_*Max*_ = *Max*(*D*_*k*_[*P*_*k*_[*i*]], *D*_*k*_[*P*_*k*_[*i* + 1]]), find range [*s, e*] such that ∀*j* ∈ [*s, e*], *D*_*k*_[*P*_*k*_[*j*]] = *D*_*Max*_, *j* ≠ *i, D*_*k*_[*P*_*k*_[*s*]] *< D*_*Max*_ and *D*_*k*_[*P*_*k*_[*e* + 1]] *< D*_*Max*_; Second, check if any *j* ∈ [*s, e*], *Y*_*k*+1_[*P*_*k*_[*j*]] = *Y*_*k*+1_[*P*_*k*_[*i*]], if so, haplotype *P*_*k*_[*i*] does not have set-maximal matches at site *k* location, and stop; Third, if it passes the second step, output a set maximal match between haplotypes *P*_*k*_[*i*] and *P*_*k*_[*j*] for all *j* ∈ [*s, e*] with a match length of *D*_*Max*_.

The scan up and scan down to find range [*s, e*] are light tasks since the scan up only has to reach the location *j*′ that *D*_*k*_[*P*_*k*_[*j*′]] ≠ *D*_*Max*_, similarly for the scan down. At this point, it is not necessary to apply complicated parallel implementation to report all versus all set-maximal matches. Instead, the parallelization is simply done by paralleling the outer loop that loops through all the haplotypes. The total work to find all the ranges [*s, e*] is *O*(*c*), where *c* is the sum of the number of haplotypes that match with haplotype *i* with length equal to the maximum match of any haplotype and *i* for all *i* ending at the current site. So, the total work among all *M* haplotypes is *O*(*M* + *c*), and the span is 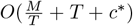 per site, where *c\** is the maximum number of haplotypes that match with haplotype *i* with length equal to the maximum match of any haplotype and *i* for all *i* ending at the current site. In practice, *c\** ≪ *M*.

## 4 Results

HP-PBWT was implemented in C# and its source code and software package are available at https://github.com/ucfcbb/HP-PBWT. To benchmark the performance of HP-PBWT, the following tests were carried out. The first test was done to ensure correctness (See Appendix B). Then, reporting all versus all L-long matches was benchmarked using HP-PBWT, Wertenbroek et al.’s parallel PBWT, and a corrected version of Durbin’s PBWT. Finally, an IO-excluded benchmark was performed for both HP-PBWT and sequential PBWT by reporting all versus all L-long matches to evaluate the scalability of the HP-PBWT-based algorithms.

### 4.1 Benchmark Design

Two sets of experiments were designed: IO-included, and IO-excluded. The IO-included experiments test real-world scenarios with hard drive input and output. VCF files were used as input for the IO-included experiments. To further evaluate the scalability of HP-PBWT, IO-excluded benchmark of HP-PBWT was implemented and an IO-excluded sequential PBWT was implemented as well. The idea of the IO-excluded benchmark was to remove the IO influence. This is similar to the run times presented by Wertenbroek et al. in the main paper. The IO-excluded benchmark is performed by randomly generating IO-excluded panels and then running the report all versus all L-long match algorithm without outputting matches to the hard drive. Dependencies were added to HP-PBWT to make sure computations were not optimized out by the compiler (See Appendix C for details).

Three measurements were used to evaluate HP-PBWT: run time, speed-up, and parallel efficiency. The run time of IO-included experiments included time for both reading the input and outputting matches. For the IO-excluded experiments, the run time was measured after generating the whole panel *X* in memory and without outputting matches. The speed-up was calculated by the sequential run time divided by the parallel run time. The parallel efficiency was calculated by speed-up divided by the number of cores.

The IO-included tests used chromosome 20 of the 1000 Genomes Project [1], and chromosome 20 of the UK Biobank [17]. To further test the scalability of these tools, randomly generated VCF files were also used with a fixed dimension of *N* = 1000 sites and *M* haplotypes that *M* = 1000 × 2^i^ where *i* ∈ [0, 15]. The reason that 1000 × 2^15^ was used as the largest input was because both Durbin’s and Wertenbroek et al.’s parallel PBWT use the same VCF handling library (HTSlib [3]) that can not read datasets with *M* ≥ 1000 × 2^16^. Thus, the largest input of these experiments was *M* = 1000 × 2^15^. The length cut-offs were 2000 sites for the 1000 Genomes Project chromosome 20 (the same as in Wertenbroek et al.’s benchmark), 1600 sites for the UK Biobank chromosome 20, and 30 sites for the randomly generated files.

The IO-excluded HP-PBWT was benchmarked against the IO-excluded sequential PBWT with *M* = 1000 ×2^i^ haplotypes for *i* ∈ [0, 31] and a fixed dimension of *N* = 100 sites. Since the *M* dimension was to reach 2 billion, only 100 sites were generated for the sake of memory capacity. The length cut-off was *L* = 50. To create stable run times all the IO-included tests were executed 10 times. For the IO-excluded tests, each experiment were executed 10 times for *i* ∈ [0, 9], 4 times for *i* ∈ [10, 17], and 2 times for *i* ∈ [18, 31].

Tests were carried out for Wertenbroek et al.’s parallel PBWT to find out which number of threads (for *T*≤ 60) had the best run time, in which when *T* = 12 Wertenbroek et al.’s parallel PBWT performed the best. This is consistent with Wertenbroek et al.’s observation. All Wertenbroek et al.’s parallel PBWT’s tests in this paper use 12 threads.

The IO-included tests were executed on local servers with Intel^R^ Xeon^R^ CPU E5-2683 v4. The IO-excluded tests were executed in Windows Server 2022 on the Amazon EC2 servers which were powered by 3.6 GHz 3rd generation AMD EPYC 7R13 processors. To eliminate the interference of different programming languages and operating systems, the IO-excluded sequential PBWT was also programmed in C#. For HP-PBWT 10, 20, 30, 40, 50, and 60 cores were used in both IO-included and IO-excluded tests, meanwhile 12 cores setting was also used in the IO-included tests, since Wertenbroek et al.’s parallel PBWT had best run time on 12 cores setting.

### 4.2 IO-Included Benchmarks

The run times in Table 1 show Wertenbroek et al.’s parallel PBWT’s best run time (from 12 threads) did not have any speed-up comparing to Durbin’s version on either chromosome 20 of UK Biobank genotyping array data or chromosome 20 of 1000 Genomes Project sequencing data. With the same 12 thread setting, HP-PBWT had 2-fold speed-up on chromosome 20 of UK Biobank genotyping array data. Meanwhile HP-PBWT’s best run time had about 4-fold speed-up on chromosome 20 of UK Biobank genotyping array data type with 30 threads. HP-PBWT did not have any speed up in 1000 Genomes Project sequencing data, since the *M* dimension of 1000 Genomes Project is too small, it only has 5008 haplotypes, and the parallelization of HP-PBWT is not designed for this type of inputs. The speed-ups show that HP-PBWT can improve the run time of PBWT in large population biobank genotyping array data. Meanwhile, further improvements are needed for HP-PBWT to deal with sequencing data with less amount of haplotypes.

**Table 1.**
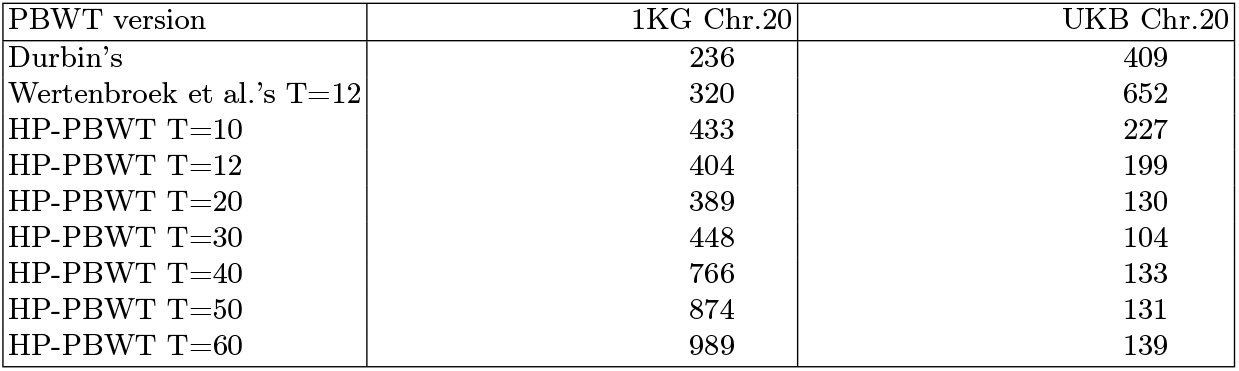
IO-included run time (in seconds) for real datasets. “1KG” stands for 1000 Genomes Project, “UKB” refers to UK Biobank.

To test HP-PBWT’s performance on large amount of population, randomly generated VCF files were used, since there was not any real dataset with the number of samples is greater than 1 million available to the authors. Figure 4(A) shows the run times of all tools increased when the input size *M* increased. The speed-up in Figure 4(B) shows the Wertenbroek et al.’s parallel PBWT’s best speed-up was about 1.6-fold, HP-PBWT had a 3-fold speed-up with 12 threads and a 7.2-fold speed-up with 60 threads on the largest input comparing to the corrected Durbin’s PBWT. It shows that HP-PBWT started to gain speed-up to the corrected Durbin’s PBWT at the input of *M* = 64*k*, meanwhile Wertenbroek et al.’s parallel PBWT started to gain speed-up at *M* = 4*m* with 1.04-fold speed-up and reached its best speed-up of 1.6-fold speed-up at *M* = 32*m*.

**Fig. 4.**
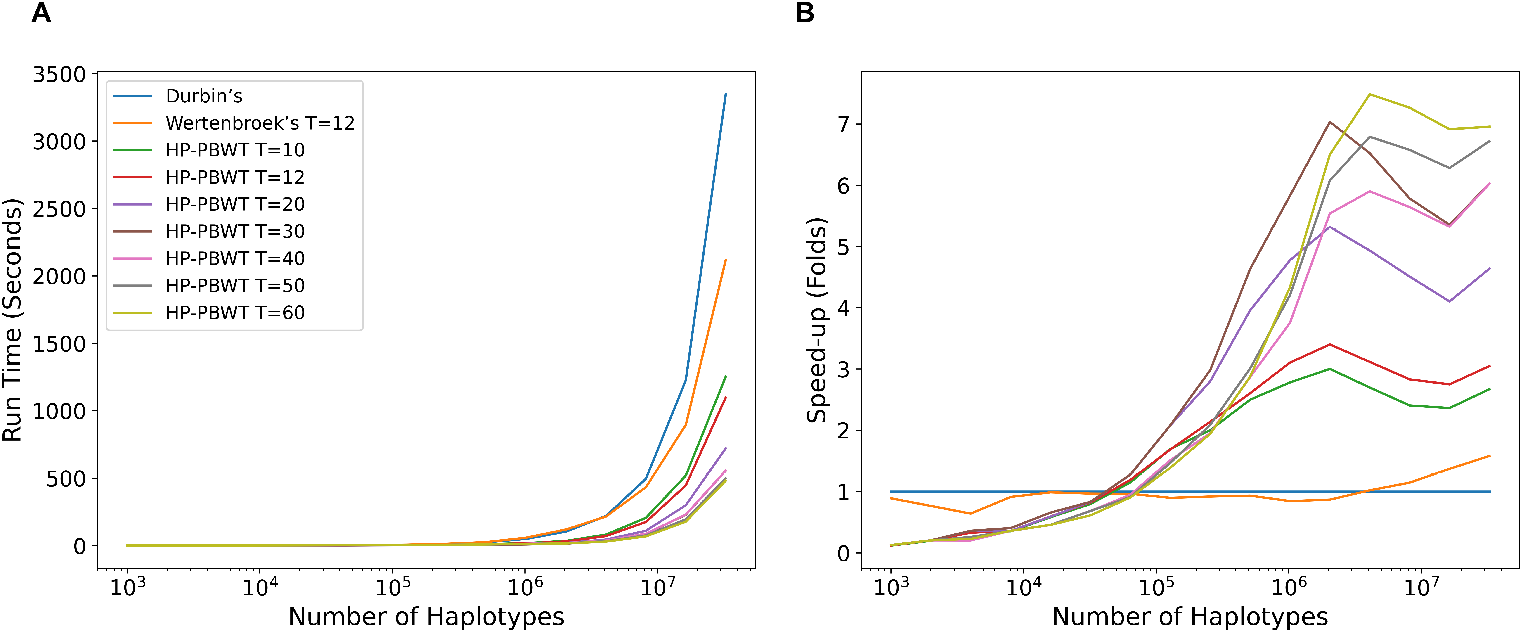
IO-included run time (A) and speed-up (B) based on the random dataset across different versions of PBWT.

### 4.3 IO-Excluded Benchmarks

Since Durbin’s PBWT does not have an IO-excluded benchmark mode and Wertenbroek et al.’s benchmark mode only reads and converts hard drive files into memory. Heavy modifications should not be applied to these versions to fit the benchmark purposes. Thus, in the IO-excluded benchmarks, the IO-excluded HP-PBWT was only benchmarked with the IO-excluded sequential PBWT.

The run time in Figure 5(A) and the speed-ups in Figure 5(B) show that HP-PBWT had better performance when the size of the input increased. The best performance of HP-PBWT against the sequential PBWT was 22.2-fold at *M* = 8, 192, 000 with 60 threads. The benchmarks also showed the maximum number of threads (60) setting used in these experiments did not have the best run time all the time. For the largest 2 billion data input, the sequential PBWT took 5.9 hours and HP-PBWT with 50 threads execution had the shortest run time, 0.4 hours, a 13.3-fold speed-up. This benchmark was performed on the Amazon EC2 servers.

**Fig. 5.**
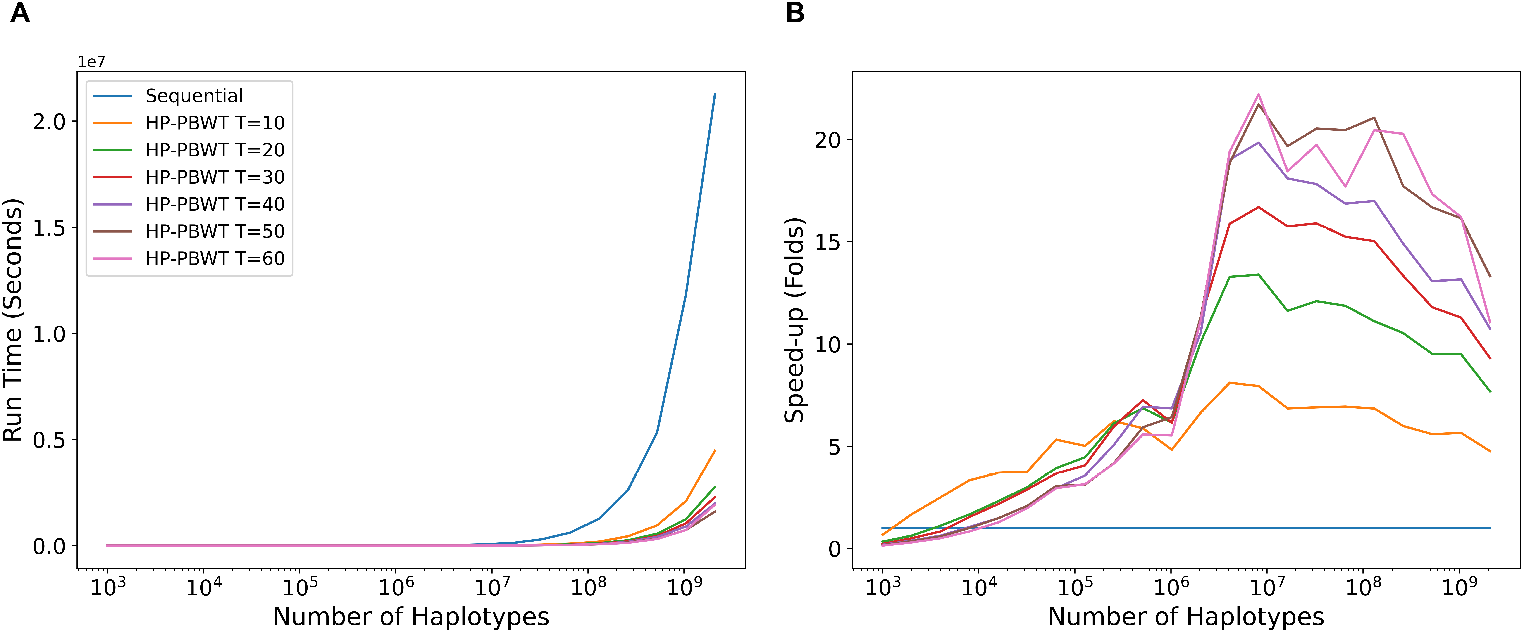
IO-excluded run time (A) and speed-up (B) based on the random dataset between HP-PBWT and reimplemented IO-excluded sequential PBWT.

The speed-up shown in Figure 5(B) (also shown in Figure 6 in Appendix D) and parallel efficiency shown in Figure 7 in Appendix D started to fall once *M* increased beyond 8 million. There are a couple of potential impact factors. First, the estimated 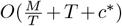 time complexity per site for reporting all versus all L-long matches is mainly dominated by the number of haplotypes *M*, but once *M* increases to a certain level *c\**, the maximum number of matches per haplotype, also becomes larger. On the other hand, this also reflects the parallel computational theory [7], that the more threads that are being used, the more idle worker time there is. Figure 8 in Appendix D shows the run time increased accordingly with the number of matches, once *M* reached to some millions the run time increased significantly. Second, CPU temperature also has a major impact on run time. That is why nowadays CPU and GPU cooling methods are being researched and developed constantly [12, 14, 15].

**Fig. 6.**
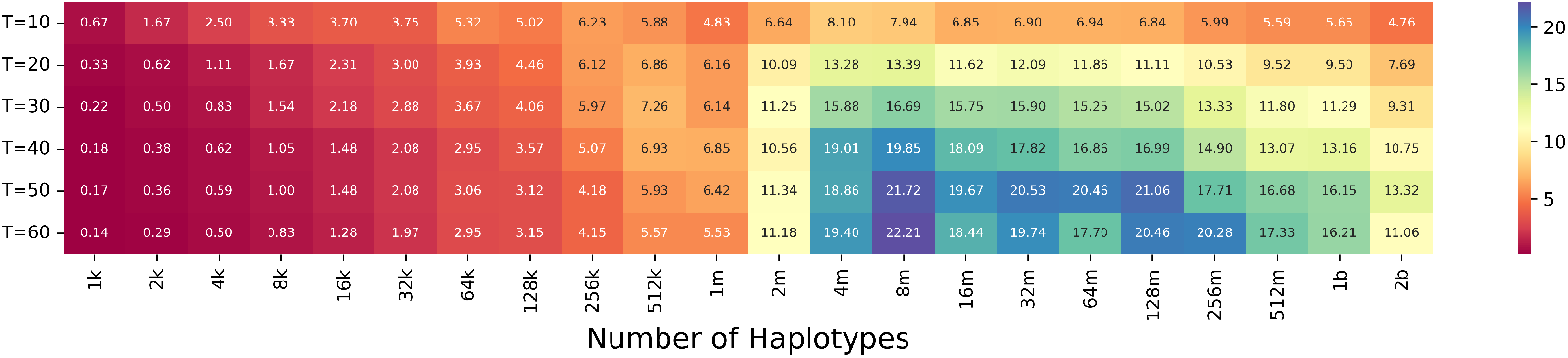
Speed-up from IO-excluded tests.

**Fig. 7.**
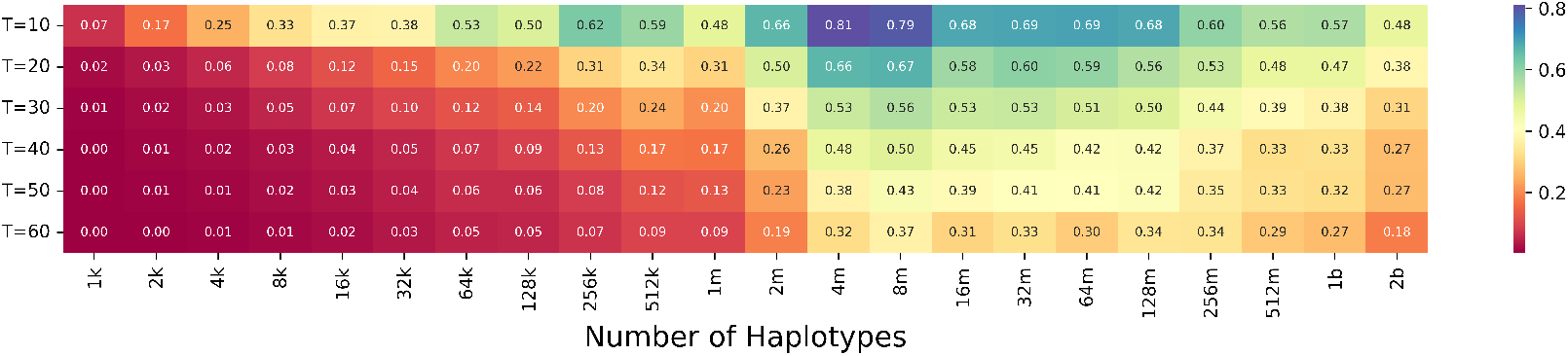
Parallel efficiency from IO-excluded tests. The parallel efficiency is calculated by speed-up divided by the number of threads.

**Fig. 8.**
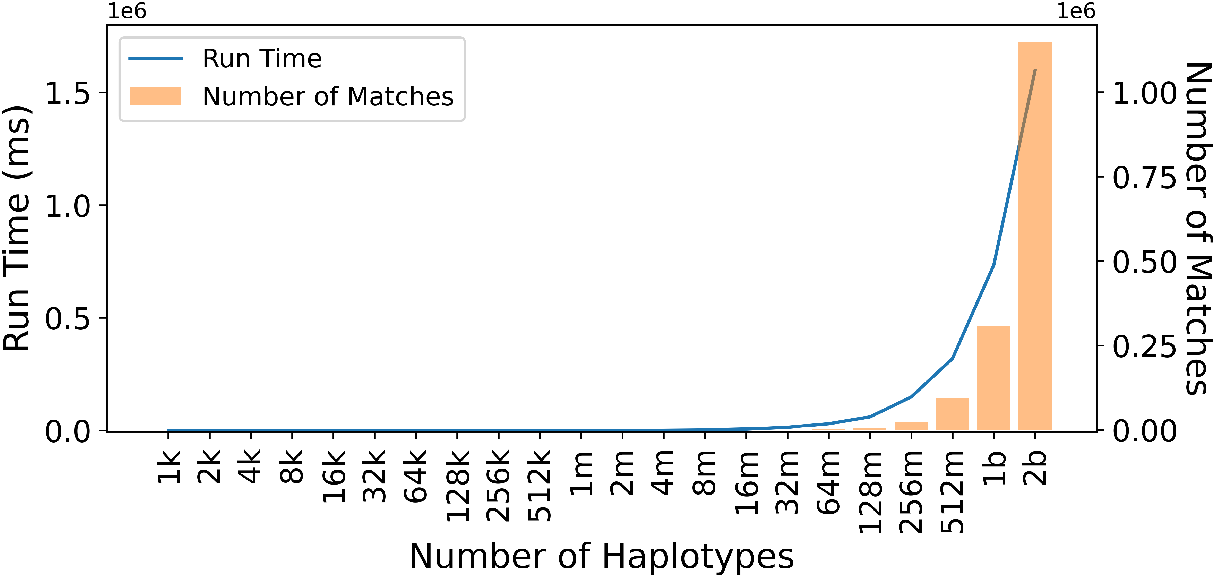
The correlation between run time (in milliseconds) and number of matches, the data was collected from IO-excluded tests with 50 threads.

## 5 Discussion

In this paper, new algorithms were designed to break the dependencies that prevent Durbin’s PBWT from being executed in parallel on the haplotype dimension, *M*. HP-PBWT algorithms enable parallel PBWT on the panel construction of prefix arrays, divergence arrays, reporting all versus all L-long matches, and reporting all versus all set-maximal matches. HP-PBWT is both memory and IO efficient. HP-PBWT can be applied on any PBWT-based applications to leverage much larger genetic panels up to billions of haplotypes efficiently.

Currently, HP-PBWT works well on biobank scale genotyping array data, but does not have speed-up on sequencing data with small amount of population. It is likely due to the partition mechanisms in prefix array and divergence array computations, they might have too much overhead. New algorithms and optimizations can be researched for the sequencing datasets. Another possible way to speed-up the whole process is to apply both haplotype-based parallel PBWT and site-based parallel PBWT algorithms simultaneously.

## Acknowledgments

This work was supported by the National Institutes of Health under award numbers R01HG010086 and R01AG081398. This research has been conducted using the UK Biobank Resource under application number 24247.

## Appendices

### A Algorithms

#### Algorithm 1

Computate of prefix array *P*_*k*_ in parallel

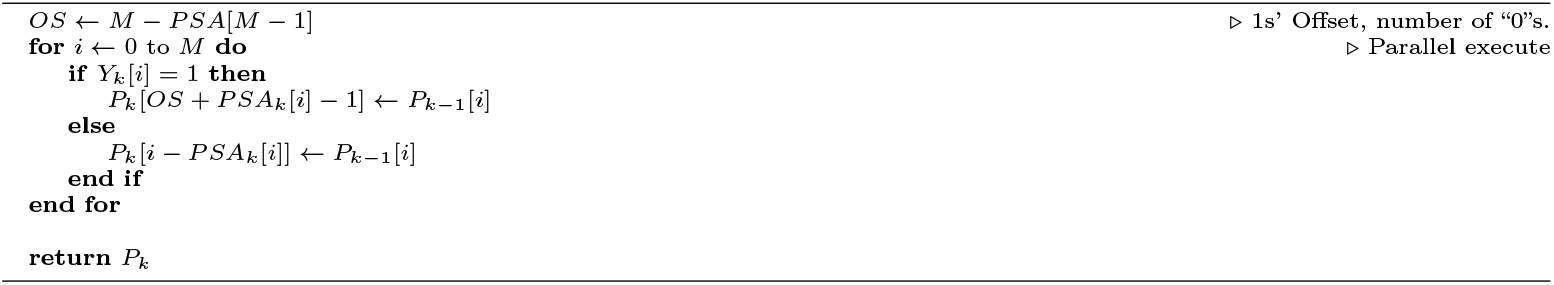

#### Algorithm 2

Compute divergence array *D*_*k*_ in parallel

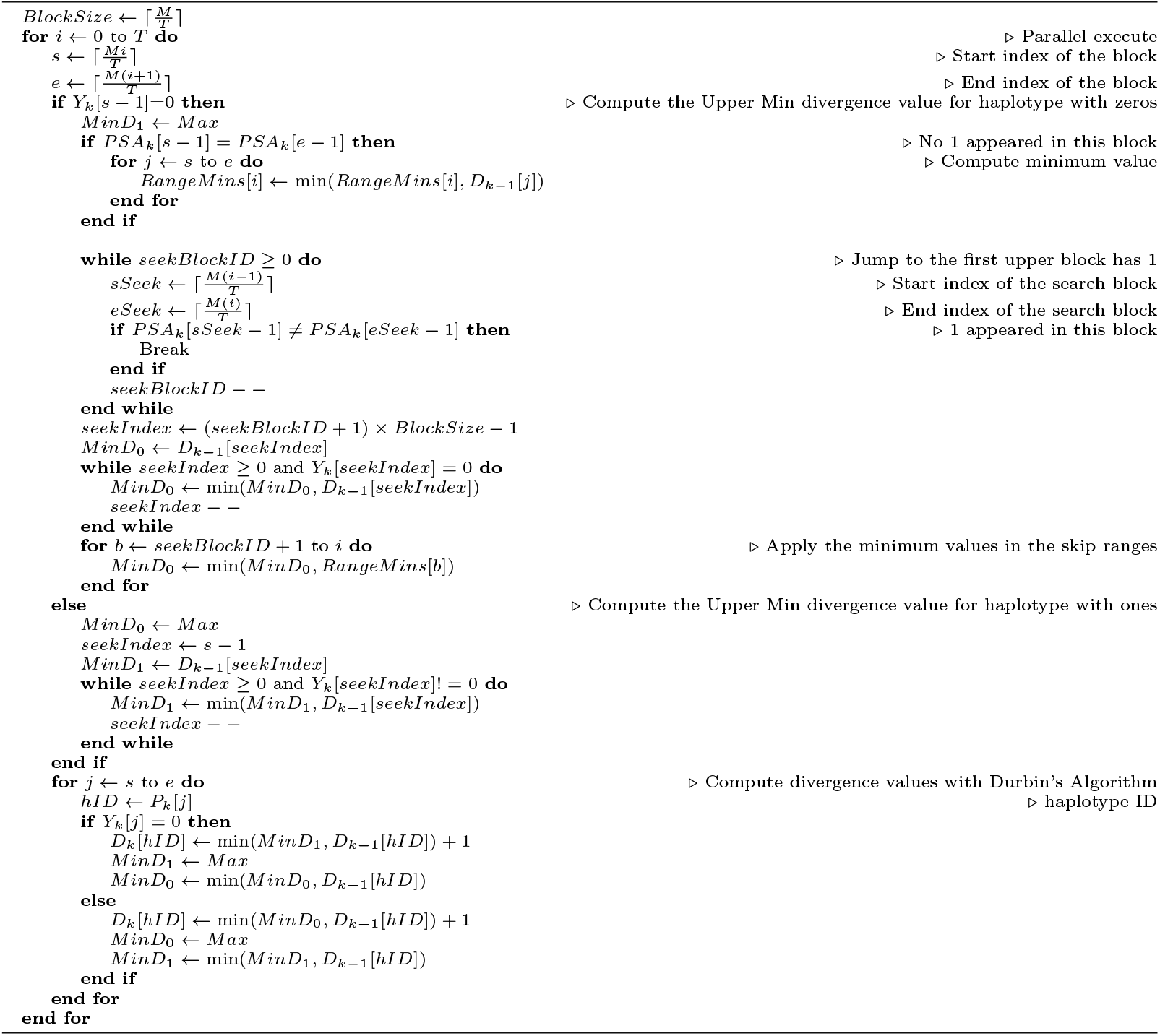

#### Algorithm 3

Report All versus All L-long matches ended at *k* site in parallel

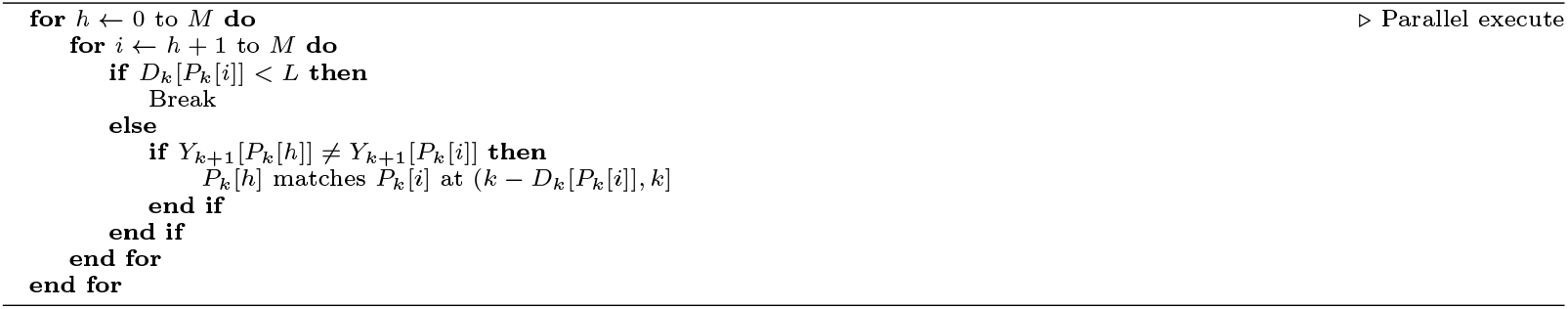

### B Correctness

To verify the correctness of HP-PBWT, first, naive algorithms that find all versus all L-long matches and all versus all set-maximal matches ended at each site were implemented. Then the matches from the IO-excluded HP-PBWT, the IO-excluded sequential PBWT, and the naive algorithms were compared to ensure the correctness. Second, HP-PBWT’s output was compared with Durbin’s PBWT, and Wertenbroek et al.’s parallel PBWT on UK Biobank chromosome 20 with *L* = 1600. Wertenbroek et al.’s parallel PBWT outputted 10,348,502 matches, Durbin’s PBWT outputted 10,362,502 matches, and HP-PBWT outputted 10,521,839 matches. Every match outputted by Wertenbroek et al’s parallel PBWT. was outputted by Durbin’s PBWT and every match outputted by Durbin’s PBWT was outputted by HP-PBWT. The matches outputted by HP-PBWT were verified, they were all versus all L-Long matches. Two minor issues were found in Durbin’s implementation. The first issue is it does not report matches that match to the last site. The second issue is the reporting code block can not be triggered when it reaches the last haplotype in *P*_*k*_ that *D*_*k*_[*P*_*k*_.*Last*] ≥*L*. After correcting these two issues, HP-PBWT and the corrected Durbin’s version produce identical outputs. Furthermore, it was observed that the corrected Durbin’s PBWT had nearly equal run time performance to the original Durbin’s PBWT. Thus, the Durbin’s run time presented in this paper are from the corrected Durbin’s PBWT.

### C Dependencies

Dependencies were added to IO-excluded HP-PBWT and the IO-excluded sequential PBWT to make sure none of the code blocks was skipped to create correct run times. The dependencies were created by maintaining a check sum for each haplotype of the haplotypes it matched to and the lengths of the matches. At the end one of the check sums was randomly selected and outputted.

### D Additional Figures

